# The consequences of polyandry for sibship structures, distributions of relationships and relatedness, and potential for inbreeding in a wild population

**DOI:** 10.1101/145987

**Authors:** Ryan R. Germain, Peter Arcese, Jane M. Reid

## Abstract

The evolutionary benefits of simultaneous polyandry (female multiple mating within a single reproductive event) remain elusive. One potential benefit could arise if polyandry alters sibship structures and consequent relationships and relatedness among females’ descendants, and thereby intrinsically reduces future inbreeding risk (the ‘indirect inbreeding avoidance hypothesis’). However such effects have not been quantified in naturally complex reproductive systems that also encompass iteroparity, overlapping generations, sequential polyandry, and polygyny. We used long-term social and genetic pedigree data from song sparrows (*Melospiza melodia*) to quantify cross-generational consequences of simultaneous polyandry for offspring sibship structures and distributions of relationships and relatedness among possible mates. Simultaneous polyandry decreased full-sibships and increased half-sibships on average, but such effects varied among females and were smaller than would occur in the absence of sequential polyandry or polygyny. Further, while simultaneous polyandry decreased the overall frequencies of possible matings among adult full-sibs, it increased the frequencies of possible matings among adult half-sibs and more distant relatives. These results imply that the intrinsic consequences of simultaneous polyandry for inbreeding risk could cause weak indirect selection on polyandry, but the magnitude and direction of such effects will depend on complex interactions with other mating system components and the form of inbreeding depression.

## Introduction

Understanding the evolutionary causes and consequences of simultaneous polyandry, defined as female multiple mating within a single reproductive event, remains a central challenge in evolutionary ecology (Arnqvist and Nilsson 2000; Jennions and Petrie 2000; Parker and Birkhead 2013; Pizzari and Wedell 2013). One key puzzle is that direct costs of multiple mating identified in diverse systems often exceed any obvious direct benefits, meaning that polyandry can decrease females’ own fitness (e.g., Rowe 1994; Fedorka et al. 2004; Cornell and Tregenza 2007; Forstmeier et al. 2014). The widespread occurrence of simultaneous polyandry consequently implies that it might provide some indirect benefit, manifested as increased fitness of polyandrous females’ descendants rather than of the polyandrous females themselves (Tregenza and Wedell 2000; Slatyer et al. 2012; Taylor et al. 2014).

Numerous potential indirect benefits of polyandry that would be manifested as increased offspring fitness have been proposed (Jennions and Petrie 2000; Slatyer et al. 2012). For instance, polyandrous females might produce female and/or male offspring of higher additive genetic or phenotypic value for fitness (e.g. Garcia-Gonzalez and Simmons 2005; Forstmeier et al. 2011; Reid and Sardell 2012), or produce offspring that are less inbred and hence express less inbreeding depression (Stockley et al. 1993; Tregenza and Wedell 2000, 2002; Michalczyk et al. 2011; Duthie et al. 2016). However, such mechanisms often require some form of active female mate choice and/or paternity allocation, which may impose additional costs such as male harassment or increased risk of predation during mate-searching (e.g., Rowe et al. 1994, 1998; Parker and Pizzari 2010; Duthie et al. 2016), or invoke genetic constraints on female strategies (Forstmeier et al. 2011, 2014). Further, empirical evidence of substantial indirect fitness benefits to polyandrous females’ offspring remains scant (Jennions and Petrie 2000; Arnqvist and Kirkpatrick 2005; Evans and Simmons 2008; Reid and Sardell 2012; Forstmeier et al. 2014; Hsu et al. 2014).

This situation raises the possibility that polyandry evolution might be facilitated by indirect benefits manifested a further generation into the future (i.e., increased fitness of polyandrous females’ grandoffspring). Indeed, the ‘indirect inbreeding avoidance hypothesis’ (IIAH, e.g., Cornell and Tregenza 2007) postulates that simultaneous polyandry directly affects the distribution of paternity among population members, and thereby alters inbreeding risk for polyandrous females’ offspring. Specifically, when polyandry causes multiple paternity, some offspring of polyandrous females are maternal half-sibs (i.e., common mother, different father), rather than full-sibs (i.e., both parents in common) as would result from monandry (fig. 1A). In situations where individuals mate locally (i.e. given restricted dispersal), polyandry might consequently reduce the potential (i.e., the expected frequency given random mating) for full-sib inbreeding among a female’s offspring (Cornell and Tregenza 2007). Grandoffspring of polyandrous females would consequently be less inbred than grandoffspring of monandrous females on average and, given inbreeding depression in fitness, contribute more offspring (i.e., great-grandoffspring of the original polyandrous female) to the population. The relative frequency of alleles underlying polyandry might consequently increase across generations. Consequently, the basic IIAH outlines a mechanism by which simultaneous polyandry could reduce inbreeding across generations, and hence facilitate its own ongoing evolution and persistence, without requiring direct inbreeding avoidance through mate choice or incurring associated costs.

**Figure 1:**
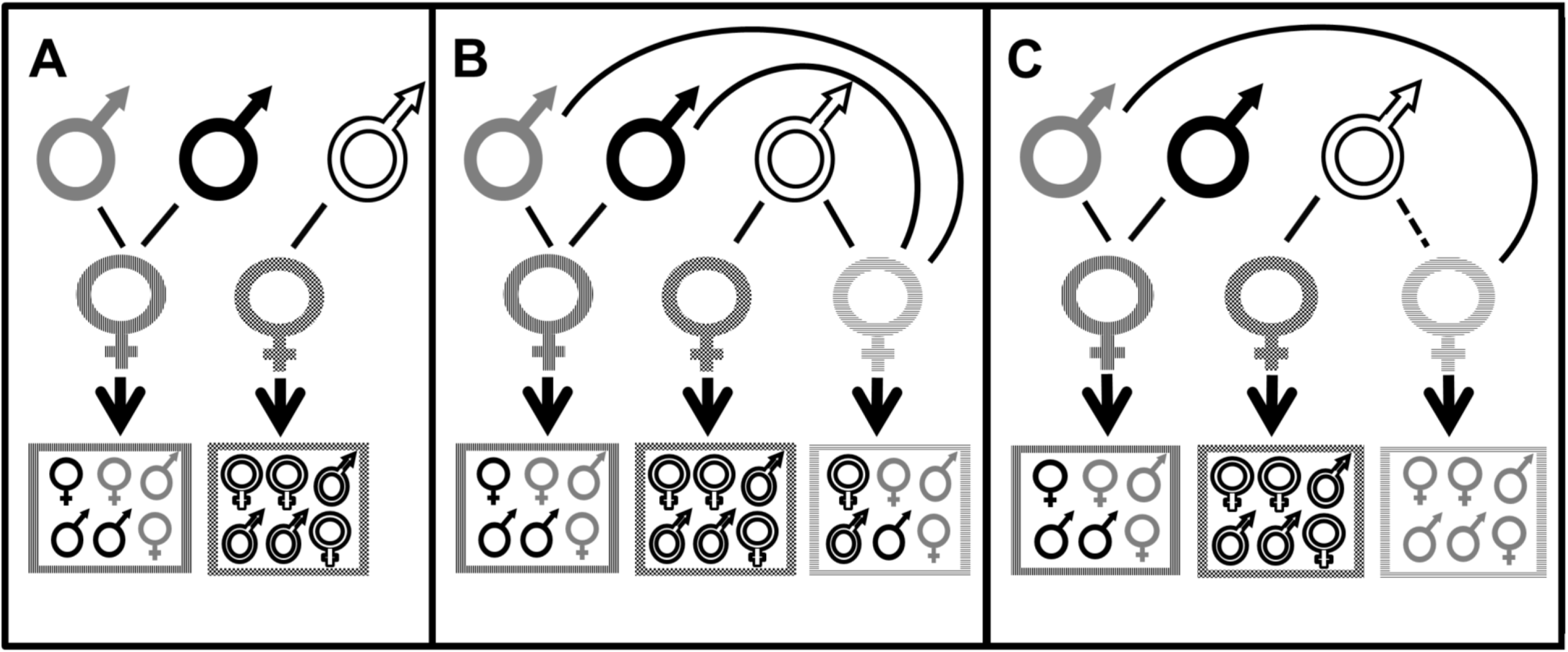
Conceptualized mating systems with simultaneous polyandry and (A) distinct males across females (‘distinct males assumption’); (B) common males across females (i.e., polygyny) with independent paternity; and (C) common males across females and nonindependent (i.e., skewed) paternity. Top female and male symbols depict breeding females and their mate(s) (connected by black lines). Boxed females and males depict resulting offspring from each mating, where box edge patterns match offspring to their mother, and individual shading match offspring to their father. In (A), a polyandrous female’s (vertical stripes) offspring have the same mother (i.e., enclosed within vertical striped box), but only some have the same father (i.e., are full-sibs rather than maternal half-sibs; matching grey or black shading). A monandrous female’s (checkered box) offspring all have the same mother and father. In (B), the same males can mate with multiple polyandrous and/or monandrous females, creating more offspring that have the same father (i.e., paternal half-sibs), and fewer unrelated offspring that share neither parent. In (C), a polyandrous female (horizontal stripes) mates with an initial male (connected by dashed line) but all of her offspring are sired by the same additional male, resulting in full-sib offspring (as for A).

To provide a first theoretical evaluation of the IIAH, Cornell and Tregenza (2007) presented a mathematical model that considers the evolutionary dynamics of polyandry resulting from reduced occurrence of full-sib inbreeding among polyandrous females’ offspring. They primarily considered the specific circumstance of non-overlapping, alternating generations of within-brood inbreeding and complete outbreeding, such as could occur in short-lived invertebrates colonizing discrete patches. Their analyses suggest that the intrinsic evolutionary benefit of the IIAH process is small, as is typical for any form of indirect selection (e.g., Kirkpatrick and Barton 1997; Møller and Alatalo 1999; Arnqvist and Kirkpatrick 2005), but might still act in combination with other benefits and appropriate genetic architecture to facilitate ongoing polyandry evolution. However, Cornell and Tregenza’s (2007) specific formulation of the IIAH makes assumptions that, while sensible in the context of their initial conceptual development and associated heuristic model, limit its direct applicability to understanding polyandry evolution in complex natural reproductive systems where polyandry and inbreeding risk co-occur.

First, Cornell and Tregenza’s (2007) formulation of the IIAH does not explicitly consider how the consequences of polyandry for the potential occurrence of inbreeding might extend beyond a polyandrous female’s immediate full-sib versus half-sib offspring and accumulate across multiple broods and generations. In iteroparous species, individuals commonly produce multiple offspring broods within and/or across years with overlapping generations. In such cases, multiple full-sibs and maternal half-sibs could be produced across different broods, for example where females mate with different initial (e.g., socially paired) males in different reproductive events (i.e., sequential polyandry) due to mate death or divorce. The set of possible mates available to a given individual offspring once they reach adulthood might then include various full-sibs and half-sibs originating from current, previous and subsequent broods produced by their mother. Moreover, it might also include full-and half-cousins and more distant full-and half-relatives, which are themselves generated across broods and generations, contingent on the degrees of simultaneous and sequential polyandry enacted by each individual’s female ancestors.

Second, Cornell and Tregenza’s (2007) formulation does not explicitly consider how the effects of polyandry on the frequencies of different relationships, and hence on the potential for different degrees of inbreeding, depend on the overall distribution of paternity within a population. Their model assumes that all polyandrous females’ additional mates are distinct, such that they do not sire offspring elsewhere in the population (hereafter the ‘distinct males assumption’). Polyandry then creates maternal half-sibs rather than full-sibs but does not create any paternal half-sibs (fig. 1A). The potential for full-sib mating among a polyandrous female’s offspring is consequently reduced, reflecting the implicit increase in effective population size. However, in many natural systems males commonly sire offspring of multiple polyandrous and/or monandrous females (i.e., polygyny, fig. 1B; e.g., Uller and Olsson 2008; Coleman and Jones 2011; Lebigre et al. 2012; McDonald et al. 2013). Such co-occurrence of polyandry and polygyny can still reduce the number of full-sibs and increase the number of maternal half-sibs compared to monandry, but can also increase the number of paternal half-sibs and reduce the number of unrelated individuals in the population (fig. 1B). Further, polyandrous females may mate with the same additional males over successive reproductive events and/or allocate all paternity to their additional mate and consequently produce more full-sibs and fewer half-sibs than otherwise expected (fig. 1C). By altering the distribution of relationships among possible mates, such paternity allocations could reduce, eliminate or even reverse the evolutionary benefit of simultaneous polyandry that the basic IIAH postulates.

Furthermore, in populations where some degree of inbreeding is common, changes in sibship structures and hence in the ‘relationships’ among possible mates resulting from polyandry may cause more complex changes in ‘relatedness’. This is because shared ancestry between a focal pair’s parents can increase the pair’s relatedness above that expected given the same immediate relationship in an outbred population. For example, the relatedness between inbred half-sibs can approach that between outbred full-sibs (Jacquard 1974; Lynch and Walsh 1998; Reid et al. 2016). Polyandry might therefore have less effect on the distribution of relatedness among possible mates than expected given its effect on the distribution of relationships.

Despite these possibilities, no studies have yet quantified the consequences of simultaneous polyandry for the distributions of sibships, relationships, and relatedness arising in natural populations. Consequently, there is no empirical basis on which to consider how the evolutionary causes and consequences of simultaneous polyandry could be influenced by the intrinsic effects of such polyandry on population-wide sibship or relationship structures and the resulting potential for inbreeding. Such investigations are particularly required for complex mating systems where iteroparity, overlapping generations, and non-independent paternity within and among females’ reproductive events can result in complex combinations of polyandry, polygyny, and mate fidelity occurring alongside inbreeding (e.g., Cockburn et al. 2003; Michalczyk et al. 2011; Culina et al. 2015, Reid et al. 2015*b*).

Effects of simultaneous polyandry on relationships and relatedness among possible mates could be quantified by experimentally enforcing polyandry or monandry across multiple generations (e.g., Power and Holman 2014). However, such experiments may simultaneously alter other life-history traits such as female fecundity or offspring survival (e.g., Fox 1993; Fedorka and Mousseau 2002; Fisher et al. 2006; Taylor et al. 2008), thereby directly altering sibship structures and relationship frequencies. Furthermore, distributions of relationships and relatedness all depend on population size and dispersal rate, on among-individual variation in survival and reproductive success, and on variation in pre-reproductive mortality of offspring sired by different males (e.g., Fisher et al. 2006; Gowaty et al. 2010; Sardell et al. 2011; Hsu et al. 2014). The composite effects of simultaneous polyandry on the potential for inbreeding could therefore be usefully quantified in free-living populations where individual reproduction and offspring survival are not artificially constrained.

One tractable approach is to utilize systems where a female’s potential and realized allocations of offspring paternity to initial versus additional mates can be documented directly. Realized distributions of relationships and relatedness emerging from realized paternity can then be compared with inferred distributions that would have emerged had all a female’s offspring in a given brood been sired by her initial mate (i.e., within-brood monandry). Socially-monogamous species with extra-pair reproduction, and hence underlying simultaneous polyandry, allow such comparisons. Here, a female’s initial socially-paired male can be identified from behavioral observations and realized paternity can be assigned by molecular genetic analysis (e.g. Webster et al. 1995, 2007; Freeman-Gallant et al. 2005; Lebigre et al. 2012). Accordingly, we used comprehensive song sparrow (*Melospiza melodia*) pedigree data to quantify the consequences of extra-pair reproduction for sibship structures and distributions of relationships and relatedness between possible mates, and thereby quantify key processes that underlie the IIAH.

First, we quantify the degree to which extra-pair reproduction changes the proportion of full-sib versus half-sib offspring produced by females over their lifetimes given realized patterns of iteroparity and social pairing and repairing, and thereby quantify the fundamental basis for the IIAH. We further quantify the degree to which observed changes differ from those predicted given lifelong monogamy and given the ‘distinct males assumption’, and thereby quantify effects of sequential polyandry and polygyny on the IIAH process. We additionally quantify how sibship structures differ among females’ hatched and adult offspring, and thereby consider the degree to which pre-reproductive mortality can shape effects of extra-pair reproduction on sibship structures among breeding individuals.

Second, we quantify the degree to which extra-pair reproduction alters the distribution of relationships among possible mates given natural iteroparity and overlapping generations, and hence alters the individual and population-wide potential for inbreeding between close and more distant relatives within the observed adult population.

Third, we quantify the degree to which extra-pair reproduction interacts with inbreeding to shape the distribution of relatedness across possible mates within and across categories of relationship. Through this sequence of three sets of analyses we elucidate the potential overall effects of the IIAH process on the population-wide potential for inbreeding in naturally complex mating systems.

## Methods

### Study system

A resident population of song sparrows inhabiting Mandarte Island, British Columbia, Canada, has been intensively studied since 1975 (Smith et al. 2006). Each year, all breeding pairs are closely monitored, all nests are located and all offspring are uniquely marked with colored plastic leg bands approximately six days after hatching (Smith et al. 2006; Wilson et al. 2007; Germain et al. 2015). Mandarte lies within a large song sparrow meta-population and receives regular immigrants (recent mean 0.9 year^-1^, ~75% female) that prevent the mean degrees of relatedness and inbreeding from increasing (Reid et al. 2006; Wolak and Reid 2016). All immigrant breeders are mist-netted and banded soon after arriving. Subsequently, the identities of all individuals alive in late April (i.e., the start of the breeding season) are recorded in a comprehensive census (resighting probability > 0.99, Wilson et al. 2007), and the socially-paired parents that rear each brood of chicks are identified (Smith et al. 2006).

Resulting data show that Mandarte’s song sparrows typically form socially monogamous breeding pairs which rear 1–4 broods of 1-4 (mean = 2.2) offspring each per year (Smith et al. 2006). Both sexes can first breed aged one year, and median adult lifespan is two years (maxima of eight and nine years in breeding females and males respectively, Smith et al. 2006; Keller et al. 2008). Due to a typically male-biased adult sex-ratio, 10–40% of males remain socially unpaired annually (Smith et al. 2006; Sardell et al. 2010; Lebigre et al. 2012). Both sexes can form new social pairings within and among years following divorce or death of their socially-paired mate (Smith et al. 2006; Reid et al. 2015*b*), and there is no sex-biased dispersal within the study system (Arcese 1989).

Extra-pair reproduction is frequent: overall, 28% of hatched offspring are sired by extra-pair males (Sardell et al. 2010; see also Hill et al. 2011), which is within the range commonly observed in passerine birds (Griffith et al. 2002). Consequently, ~45% of broods show mixed paternity, while ~10% of broods contain ≥ 2 offspring that are all sired by the same extra-pair male. Population-wide extra-pair paternity is distributed across multiple males rather than monopolized by few males (Reid et al. 2011a; Lebigre et al. 2012; Reid and Sardell 2012).

Overall, this system has proved valuable for understanding variation in mating strategy and fitness occurring in natural viscous meta-populations (i.e. with restricted dispersal) where relatives and non-relatives interact. Specifically, previous analyses showed substantial opportunity for inbreeding and inbreeding avoidance, but little evidence of active inbreeding avoidance through non-random social pairing (Keller and Arcese 1998; Reid et al. 2006) or non-random extra-pair reproduction (Reid et al. 2015*a*,*b*) with less closely related mates, despite strong inbreeding depression in fitness (Keller 1998; Reid et al. 2014; Nietlisbach et al. 2017). Further, female extra-pair reproduction is heritable (Reid et al. 2011b) but females receive no obvious direct benefits (e.g., nuptial gifts, offspring provisioning) from extra-pair males, and extra-pair reproduction can reduce offspring fitness (Sardell et al. 2012; Reid and Sardell 2012). However, the potential role of the IIAH process in maintaining extra-pair reproduction, and underlying simultaneous polyandry, has not previously been examined.

### Social and genetic pedigrees

Fully evaluating the IIAH process requires quantifying sibship structures, relationships and relatedness, which can all be calculated from pedigree data linking offspring to parents. We first compiled a ‘social pedigree’ linking all banded offspring to their observed mother and her socially-paired male spanning 1975–2015 (Reid et al. 2014, 2015a,b). Since 1993, all adults and banded offspring were blood sampled and genotyped at ~160 highly polymorphic microsatellite loci, and all offspring were assigned to genetic sires with >99% individual-level statistical confidence (Nietlisbach et al. 2015, 2017; Reid et al. 2015*a*). We then compiled a ‘genetic pedigree’ linking all banded offspring to their mother and true genetic father (Sardell et al. 2010; Reid et al. 2014, 2015*a*, 2015*b*; Nietlisbach et al. 2015). We thereby generated two parallel pedigrees spanning 1993–2015 that describe sibship structures and the distributions of relationships and relatedness among all population members as they would have been had all observed breeding pairs been monogamous within broods (‘social pedigree’), and given the realized pattern of extra-pair reproduction and underlying polyandry (‘genetic pedigree’, Lebigre et al. 2012; Reid et al. 2014). Because there is no extra-pair maternity (Sardell et al. 2010), the two pedigrees differ only in the paternity of ~28% of individuals, and are identical in terms of individual longevity, female reproductive success, and offspring survival to recruitment. Differences in sibship structure, relationships and relatedness among possible mates between the two pedigrees therefore stem solely from extra-pair reproduction (see Discussion).

To maximize use of all available pedigree data and relax the alternative assumption that all 1993 breeders are unrelated, we grafted each of the 1993–2015 social and genetic pedigrees onto the basal 1975–1992 social pedigree (Reid et al. 2014, 2015*a*). To minimize error in estimates of relationships and relatedness stemming from inadequate pedigree depth and/or remaining paternity error for some individuals hatched during 1975–1992, we restricted analyses to adults alive during 2008–2015. All such individuals had genetically-verified ancestors back to all great-great-grandparents, or were descendants of immigrants, meaning that any error due to misassigned paternities before 1993 was trivial (Reid et al. 2015*a*). Immigrants are assumed to be unrelated to existing residents, and therefore to all possible mates, in their arrival year (Marr et al. 2002, Reid et al. 2006, 2014, 2015*a*), and this assumption is supported by comparisons among neutral microsatellite marker data (Keller et al. 2001; Nietlisbach et al, unpublished data). However, immigrants could potentially inbreed with their own descendants in subsequent years.

### Sibship structures

To quantify the degree to which extra-pair reproduction altered the proportions of full-sibs versus half-sibs that each female produced over her lifetime, we compared sibship structures between the social and genetic pedigrees. We first calculated each female’s total lifetime number of banded offspring (*j*) and calculated the total number of sibships (i.e., all possible full-sib and half-sib relationships, hereafter *N*_sibs_) among the *j* offspring as 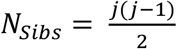. We then calculated the numbers of full-sibships and maternal half-sibships among each female’s offspring given the social and genetic pedigrees, and divided these numbers by *N*_sibs_ to obtain the lifetime proportions of full-sibships (Prop_Full-sibs_) and half-sibships (Prop_Half-sibs_) produced by each female (where Prop_Half-sibs_ = 1 - Prop_Full-sibs_) given each pedigree. The absolute difference between each female’s value of Prop_Full-sibs_ given the social and genetic pedigrees (i.e., Diff_social-gen_ = |Prop_Full-sibs[social]_ - Prop_Full-sibs[genetic]_|) quantifies the effect of extra-pair reproduction (i.e., simultaneous polyandry) on sibship structures while fully accounting for natural patterns of variation in paternity stemming from female re-pairing between broods (i.e., sequential polyandry) and repeat mating with the same extra-pair male across multiple broods.

We then undertook analyses to explicitly quantify the combined effects of simultaneous and sequential polyandry, and of polygyny, on sibship structures. First, we quantified the difference between Prop_Full-sibs_ given the genetic pedigree and the value of Prop_Full-sibs_ that would arise given strict lifelong monandry (i.e., 1.0, hence Diff_life_monandry-gen_ = 1.0 - Prop_Full-sibs[genetic]_). Second, to quantify the degree to which observed sibship structures differed from those that would have arisen in the absence of polygyny (i.e. given the ‘distinct males assumption’ that is implicit in the basic IIAH, see Introduction), we additionally considered a hypothetical pedigree in which extra-pair males could sire multiple offspring within a given brood but could not sire other within-pair or extra-pair offspring in the population (i.e., fig. 1A). We assigned a unique sire identity to all extra-pair offspring in each observed brood, maintaining the observed paternity distribution (i.e., *X_i_* extra-pair offspring sired by male *i*), and then recalculated Prop_Full-sibs_ and Prop_Half-sibs_ for each female. Finally, to elucidate mechanisms underlying observed changes in sibship structures, we also calculated the total number of males that sired at least one of each female’s offspring given the social, genetic, and ‘distinct males’ pedigrees.

We fitted generalized linear mixed models (GLMMs) to test whether the sibship structures of females’ offspring (binomial error structures, with Prop_Full-sibs_ and *N*_sibs_ as the binomial numerator and denominator, respectively), or the number of different sires (Poisson error structures), differed between the pedigrees. These models included fixed effects of pedigree (three levels) and random female identity effects. Goodness of fit (*R^2^*) for each model was assessed by the conditional coefficient of determination (Nakagawa and Schielzeth 2013). We used Tukey’s post-hoc tests to evaluate pairwise differences in Prop_Full-sibs_ and number of sires among the three pedigrees at *α* = 0.05. To quantify how differences in sibship structure varied with the degree to which individual females expressed extra-pair reproduction, we fitted further generalized linear models (GLMs) to quantify how Diff_social-gen_ and Diff_life_monandry-gen_ varied with whether or not any of a female’s offspring were sired by an extra-pair male (Supporting Information S1), or with the overall proportion of their lifetime offspring that were sired by an extra-pair male. These GLMs had binomial error structures, with Diff_social-gen_ and Diff_life_monandry-gen_ as respective binomial numerators, and *N*_sibs_ as the binomial denominator. There was little over-dispersion in our dataset beyond that accounted for by the fitted models.

All the above analyses were implemented across each female’s offspring that survived to banding, and across offspring that survived to age one year (recruits). These two sets of analyses respectively elucidate the direct primary effects of the distribution of paternity on sibship structures, and elucidate the net effects of this distribution coupled with pre-reproductive mortality on realized sibship structures among (potentially) reproductive adults. Females that were still alive in 2016, or that produced ≤1 banded or ≤1 recruited offspring (meaning that *N*_sibs_ = 0), were excluded from the respective analyses. Analyses for banded offspring were also repeated across the subset of females that produced ≥2 recruited offspring, thereby allowing direct comparison across offspring stages within females (Supporting Information S2). While our primary analyses focused on sibship structures among females’ offspring, further analyses demonstrated similar structures among males’ offspring (Supporting Information S3).

### Distribution of relationships among possible mates

We next quantified how changes in sibship structures resulting from extra-pair reproduction translated into cross-generational differences in relationships among possible mates within the observed adult population, and hence affected the potential for inbreeding. We used annual censuses of all adults alive in each year during 2008–2015 (annual means of 26.9±8.8SD females [range 13–38] and 35.1±10.5SD males [range 20–56]) to generate all possible female-male pairs that could possibly have mated in each year, assuming no mating constraints (hereafter ‘all possible matings’). Since we analyzed relationships from the female perspective the assumption of no constraints is reasonable; due to extra-pair reproduction any adult female could possibly mate with any adult male in the population.

We compared the frequencies of all possible matings for each adult female in each year that comprised key relationships given the social and genetic pedigrees. These relationships comprised: fathers, full-brothers, and sons (1^st^ degree relatives); grandfathers, uncles, half-brothers, double first cousins (i.e., both parents of each mating individual are full-sibs), nephews, and grandsons (2^nd^ degree relatives); and great-grandfathers, single first cousins, and great-grandsons (3^rd^ degree relatives). We also considered half-uncles, half-single first cousins (i.e., one parent of each mating individual is a half-sib), and half-nephews (4^th^ degree relatives) and thereby quantified effects of extra-pair reproduction (and consequent production of half-sibs rather than full-sibs) on possible matings that would otherwise have involved 3^rd^ degree relatives. Matings involving immigrants were defined as ‘unrelated’ except where immigrants could mate with their own descendants. All possible matings that did not fall into any of the above categories were considered ‘more distantly related’. If a possible mating fell into multiple categories (e.g., one case where a possible mate was both a female’s son and grandson [i.e., the progeny of a female mating with another son]) it was allocated to the closer relationship. These full- and half-relationships provide a mechanistic link between the generation of half-sibs caused by polyandry, and resulting cross-sex relationships among possible mates.

We used Wilcoxon matched pair tests to evaluate whether the lifetime number of possible matings between individual adult females and available adult males in each relationship category differed between the genetic versus social pedigrees. While changes in some relationships given the genetic pedigree may be counted in multiple years (if both the female and possible mate survive across years), these represent separate potential opportunities for inbreeding given random mating, and were thus retained. Although each female has exactly one father in each pedigree, changes in assigned father between the two pedigrees could change whether or not a female’s assigned father is still alive in certain years and hence available as a possible mate. Since there is no extra-pair maternity, the number of possible female-son matings cannot change between the two pedigrees. However, such matings were counted to provide a complete summary of possible matings among 1^st^ degree relatives.

### Distribution of relatedness among possible mates

Given the occurrence of ancestral inbreeding in a population, sibship structures resulting from polyandry, and consequent frequencies of relationships between possible mates, do not translate directly into fixed degrees of relatedness. Hence, to quantify how polyandry translates into quantitative differences in relatedness among possible mates, we used standard pedigree algorithms (Lange 1997) to calculate the coefficient of kinship (*k*) between all adult females and all available adult males given the social (*k_SOC_*) and genetic (*k_GEN_*) pedigrees. The coefficient *k* measures the probability that two homologous alleles sampled from two individuals will be identical by descent relative to the pedigree baseline, and equals the coefficient of inbreeding (*f*) of resulting offspring (Jacquard 1974; Lynch and Walsh 1998; Reid et al. 2016).

We quantified differences in *k* between each individual adult female and her lifetime set of possible mates given the genetic and social pedigrees in three ways. First, to retain the mechanistic links with relationships and underlying sibship structures, we quantified the differences in each female’s mean *k_SOC_* and *k_GEN_* with all possible mates that were identified as 1^st^, 2^nd^, 3^rd^, and 4^th^ degree relatives, or as more distantly related or unrelated, given the social pedigree. Second, to quantify the effect of extra-pair production on the *k* between each possible female-male pair, we calculated the difference in *k* for each possible mating as *k_DIFF_* = *k_GEN_ - k_SOC_*, calculated mean *k_DIFF_* for each individual female, and quantified the proportion of females for whom mean *k_DIFF_* increased, decreased, or did not change given the genetic versus social pedigrees. Finally, we quantified the degree to which extra-pair reproduction altered the overall potential for inbreeding across the whole population. To do so, we pooled all possible matings during 2008–2015 and used a two-sample Anderson-Darling test to test whether the shapes of the continuous distributions of *k_GEN_* and *k_SOC_* differed significantly (using 10,000 resampling permutations).

Analyses were run in R version 3.2.2 (R Development Core Team 2015) using packages *MasterBayes, nadiv, lme4, kinship2*, and *kSamples* (Hadfield et al. 2006; Wolak 2012; Sinnwell et al. 2014; Bates et al. 2015; Scholz and Zhu 2015). Raw means are reported ±1SD.

## Results

### Sibship structure of banded offspring

A total of 98 female song sparrows alive during 2008–2015 produced at least two banded offspring over their lifetime (mean 11.4±10.6; median 7–8, range 2–60), and hence at least one sibship. Table 1A,B summarizes the number of sires and Prop_Full-sibs_ among these females’ lifetime banded offspring given the social, genetic and ‘distinct males’ pedigrees.

**Table 1:** Summary statistics (left panel) and generalized linear mixed models (right panel) estimating differences in the number of males that sired a female’s offspring, and the proportion of full-sibships (Prop_Full-sibs_) among females’ offspring given the social, genetic and ‘distinct males’ pedigrees. Focal females and offspring comprise (A and B) banded offspring of all females that produced ≥2 banded offspring (i.e., ≥1 sibship, *n* =98 females), and (C and D) banded offspring and (E and F) recruited offspring of females that produced ≥2 recruited offspring (*n* =37 females). Raw means are presented ± 1 standard deviation (SD). Models assumed (A,C,E) Poisson or (B,D,F) binomial error structures. Estimated pedigree effects (on latent scales) are differences from the intercept (social pedigree) and are presented ±1 standard error (SE), *df* is the residual degrees of freedom, *R*^*2*^ is the conditional coefficient of determination, and *Z* and *p* values are presented for each fixed effect level where the social pedigree represents the intercept. ‘Tukey’ summarizes a Tukey post-hoc test assessing differences among pedigrees, where different lower case letters (a,b,c) represent groups with significantly different means.

| Response variable | Pedigree | Mean (±SD) | Median (Range) | df | R<sup>2</sup> | Estimate (±SE) | Z | p | Tukey |
| --- | --- | --- | --- | --- | --- | --- | --- | --- | --- |
| Banded offspring – Full Dataset |  |  |  |  |  |  |  |  |  |
| A) Number of males | Social | 1.9 (1.2) | 1 (1–6) | 290 | 0.52 | 0.45 (0.10) |  |  | a |
|  | Genetic | 2.9 (2.1) | 2 (1–10) |  |  | 0.45 (0.09) | 4.8 | <0.001 | b,c |
|  | Distinct males | 3.5 (2.9) | 2 (1–14) |  |  | 0.63 (0.09) | 7.0 | <0.001 | c |
| B) Prop<sub>Full-sibs</sub> | Social | 0.74 (0.31) | 1.00 (0.15–1.00) | 290 | 0.61 | 1.36 (0.24) |  |  | a |
|  | Genetic | 0.53 (0.34) | 0.40 (0.00–1.00) |  |  | -0.58 (0.03) | 18.6 | <0.001 | b |
|  | Distinct males | 0.49 (0.35) | 0.33 (0.00–1.00) |  |  | -0.77 (0.03) | 24.1 | <0.001 | c |
| Banded offspring – Restricted dataset |  |  |  |  |  |  |  |  |  |
| C) Number of males | Social | 2.7 (1.5) | 2 (1–6) | 107 | 0.56 | 0.86 (0.13) |  |  | a |
|  | Genetic | 4.3 (2.5) | 4(1–10) |  |  | 0.48 (0.13) | 3.8 | <0.001 | b,c |
|  | Distinct males | 5.2(3.2) | 5 (1–14) |  |  | 0.68 (0.12) | 5.6 | <0.001 | c |
| D) Prop<sub>Full-sibs</sub> | Social | 0.56 (0.32) | 0.46 (0.15–1.00) | 107 | 0.33 | 0.08(0.20) |  |  | a |
|  | Genetic | 0.40 (0.28) | 0.30 (0.10–1.00) |  |  | -0.52 (0.03) | 15.1 | <0.001 | b |
|  | Distinct males | 0.34 (0.25) | 0.25 (0.05–1.00) |  |  | -0.66 (0.04) | 19.0 | <0.001 | c |
| Recruited offspring |  |  |  |  |  |  |  |  |  |
| E) Number of males | Social | 1.6 (0.6) | 2 (1–3) | 107 | 0.18 | 0.43 (0.14) |  |  | a |
|  | Genetic | 2.2 (1.2) | 2 (1–7) |  |  | 0.31 (0.17) | 1.9 | 0.06 | a |
|  | Distinct males | 2.3 (1.3) | 2 (1–7) |  |  | 0.37 (0.17) | 2.2 | 0.03 | a |
| F) Prop<sub>Full-sibs</sub> | Social | 0.67 (0.37) | 0.85 (0.00–1.00) | 107 | 0.37 | 0.82 (0.27) |  |  | a |
|  | Genetic | 0.47 (0.37) | 0.40 (0.00–1.00) |  |  | -0.96 (0.14) | 7.0 | <0.001 | b,c |
|  | Distinct males | 0.43 (0.38) | 0.33 (0.00–1.00) |  |  | -1.06 (0.14) | 7.6 | <0.001 | c |

Given the social pedigree, the mean number of sires per female was 1.9, and mean Prop_Full-sibs_ was 0.74 (table 1A,B; fig. 2A,B). Thus, even without considering extra-pair reproduction (i.e., simultaneous polyandry), the occurrence of re-pairing between breeding events (i.e., sequential polyandry), meant that the mean proportion of full-sibships among females’ banded offspring was on average ~26% less than expected under lifelong monandry (i.e., 1.0).

**Figure 2:**
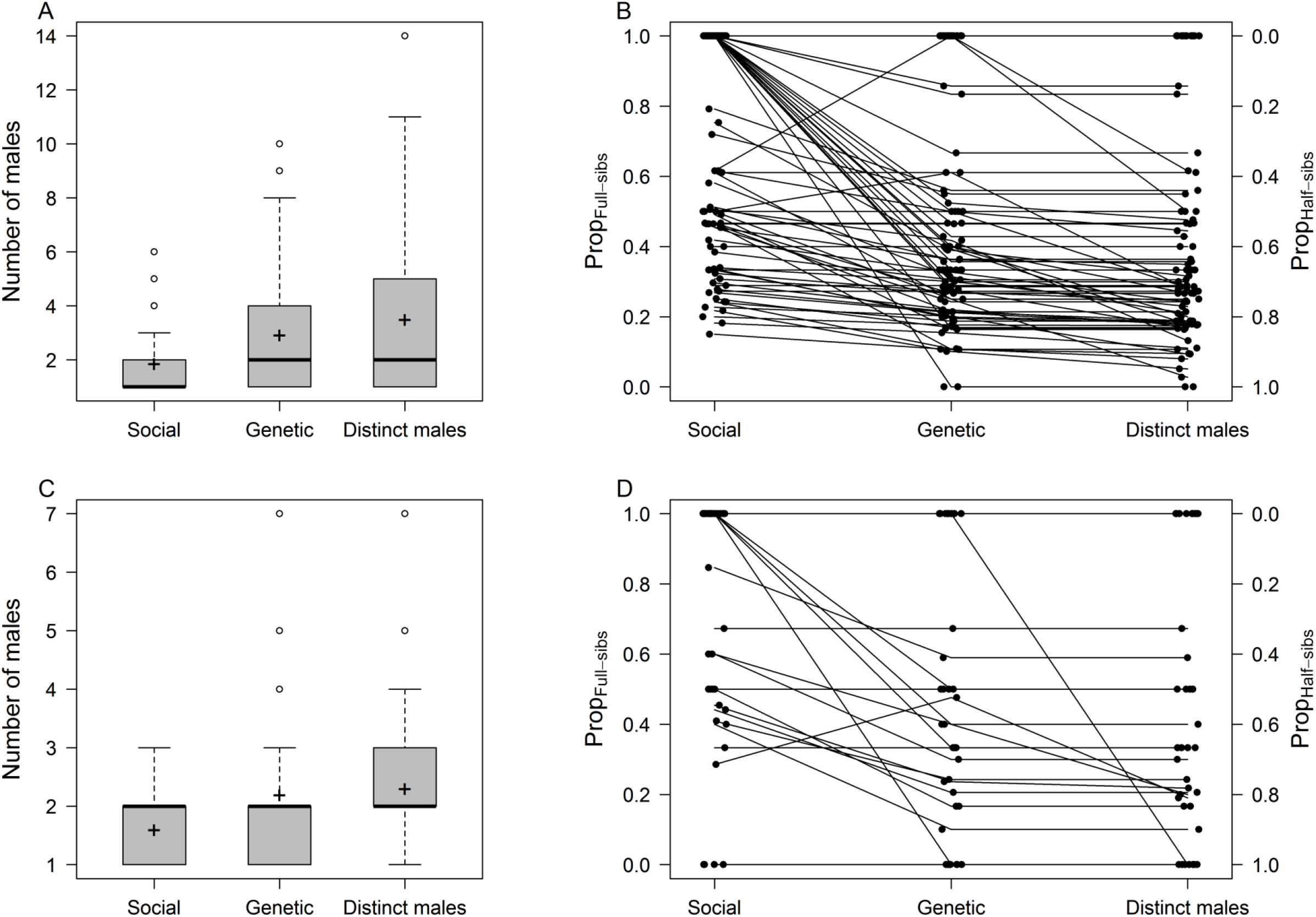
The numbers of different males that sired female song sparrows’ (A) banded and (C) recruited offspring, and the sibship structures of females’ (B) banded and (D) recruited offspring given the social pedigree (‘Social’), genetic pedigree (‘Genetic’), and ‘distinct males’ pedigree (‘Distinct males’). In (A) and (C), box lines represent the median, upper and lower quartiles, whiskers demarcate 1.5× the interquartile range, and ‘+’ shows the mean. In (B) and (D), the left and right axes respectively show the proportions of sibships among each female’s offspring that are full-sibships (Prop_Full-sibs_) and half-sibships (Prop_Half-sibs_), where points denote individual females (jittered for clarity), and lines join observations for individual females given the three pedigrees.

Given the genetic pedigree, the mean number of sires per female was 2.9, equating to a mean increase of 1.0 sire per female compared to the social pedigree (table 1A; fig. 2A). Consequently, as might be expected, Prop_Full-sibs_ among the banded offspring of most females (60%; 59/98) was lower given the genetic pedigree than given the social pedigree (table 1B; fig. 2B). However, for 38% (37/98) of females there was no change, and 2% (2/98) of females actually had higher Prop_Full-sibs_ given the genetic pedigree, illustrating that polyandry can increase rather than decrease full sibships (fig. 2B). Indeed, mean Diff_social-gen_ was greater in females where at least one offspring was sired by an extra-pair male, but greatest in females with intermediate proportions of extra-pair offspring (fig. 3A; Supporting Information S1). However, the realized effects of extra-pair reproduction on sibship structure (i.e., Diff_social-gen_ fig. 3A) were smaller, due to sequential polyandry, than would be observed had all females been strictly monandrous throughout their lifetimes (i.e., Diff_life_monandry-gen_, fig. 3B).

**Figure 3:**
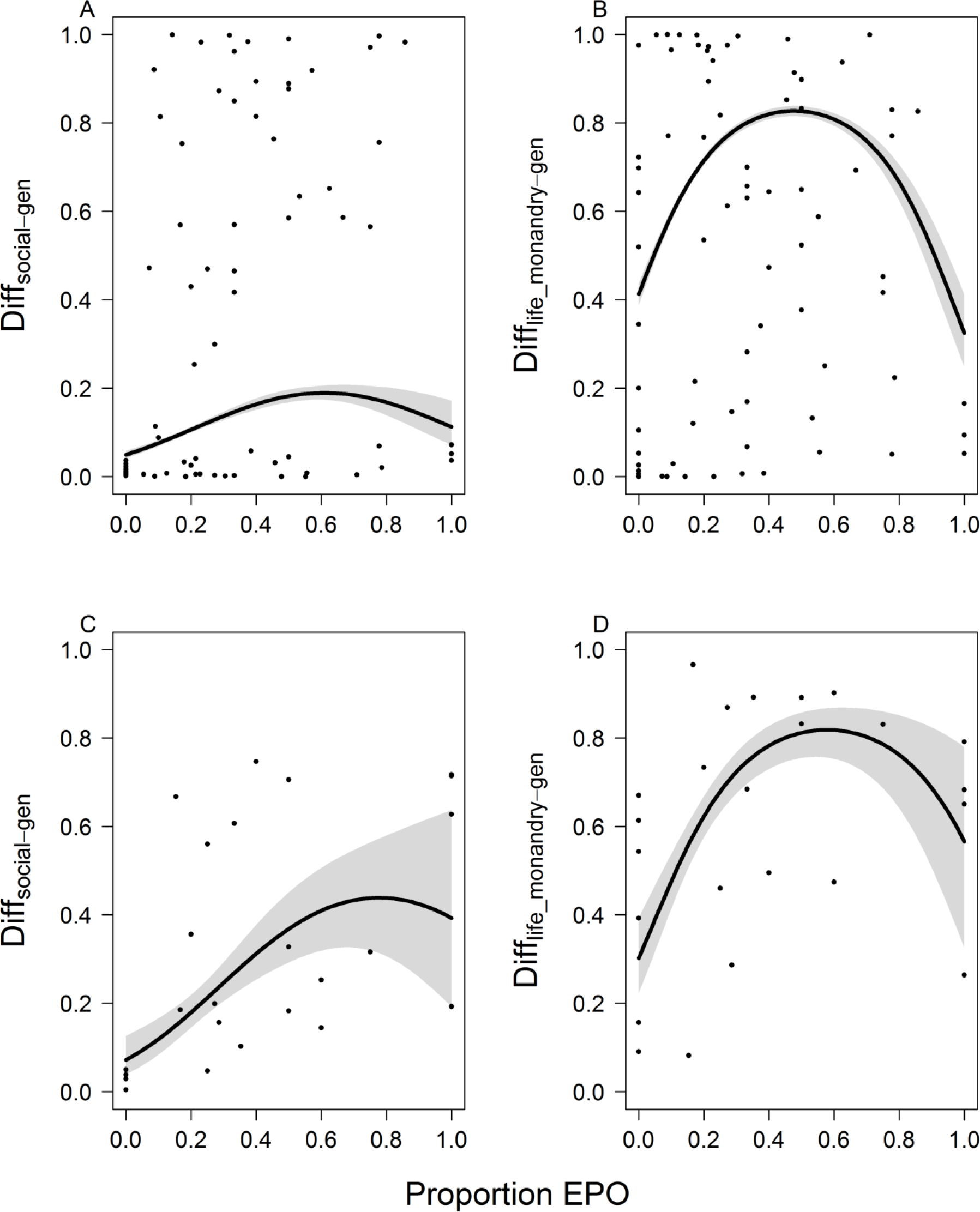
Relationships between the proportion of a female’s lifetime offspring that were extra-pair offspring (Proportion EPO) and the absolute difference in Prop_Full-sibs_ given (A and C) the genetic versus social pedigrees (Diff_social-gen_), and (B and D) the genetic pedigree versus strict lifelong monandry (Diff_life_monandry-gen_) for (A and B) banded and (C and D) recruited offspring. Predictions (black lines) and confidence intervals (grey bands) are from generalized linear models (Supporting Information S1).

As expected, the number of sires per female was greatest given the ‘distinct males’ pedigree (fig. 2A), but in fact did not differ significantly from the genetic pedigree (table 1A). However, most females (69%, 68/98) had even lower Prop_Full-sibs_ given the ‘distinct males’ pedigree than given the genetic pedigree and no females had higher Prop_Full-sibs_ (fig. 2B), creating a mean reduction in Prop_Full-sibs_ of ~8% relative to the genetic pedigree (table 1B). Thus, while female song sparrows would produce offspring with similar numbers of males given the ‘distinct males assumption’ as in reality (i.e., given the genetic pedigree), they would produce fewer full-sibships.

### Sibship structure of recruited offspring

A total of 37 females produced at least two recruited offspring over their lifetime (mean 4.2±3.0; median 3, range 2–13). Across these females, the numbers of males that sired banded offspring was consistently higher and Prop_Full-sibs_ were consistently lower than across the full set of 98 females (table 1A,B vs C,D). This is because females that produced ≥2 recruits typically produced numerous banded offspring spanning multiple broods. However, the patterns of differences between the pedigrees mirrored those estimated across all 98 females (Tukey tests, table 1A,B vs C,D).

Comparisons within the 37 females showed that the mean number of sires decreased between banded and recruited offspring, as might be expected given offspring mortality, and hence no longer differed as substantially among the three pedigrees (table 1C,E). Meanwhile, mean Prop_Full-sibs_ was slightly higher for recruited offspring than for banded offspring across all three pedigrees (table 1D,F), but mean Prop_Full-sibs_ among recruited offspring was again lower given the genetic versus social pedigrees (table 1F). At the individual level, 46% (17/37) of females had lower Prop_Full-sibs_ given the genetic pedigree, while 51% (19/37) had no change and one female had higher Prop_Full-sibs_ (fig. 2D). Diff_social-gen_ was again greater in females with intermediate proportions of extra-pair offspring (fig. 3C), and the effects of extra-pair reproduction on recruit sibship structure were smaller than would be observed given lifelong monandry (fig. 3D, Supporting Information S1). Finally, the difference in Prop_Full-sibs_ given the genetic versus ‘distinct males’ pedigrees was no longer significant across recruited offspring (Tukey test, table 1F; fig. 2D). Thus, while patterns in the effects of extra-pair mating on sibship structure were qualitatively similar among banded and recruited offspring, these effects were more pronounced among banded offspring, suggesting that early offspring mortality can reduce or alter the effects of polyandry on sibship structures.

### Distribution of relationships among possible mates

There was a total of 8028 possible matings between adult females and adult males that were alive in each year during 2008–2015, spanning 114 females and 144 males. On average, there were 0.6 fewer possible matings between individual focal females and their full-brothers given the genetic versus social pedigrees, but 1.6 more possible matings with half-brothers (table 2). However the distributions of the within-female differences in the numbers of full- and half-brothers between the two pedigrees spanned zero, showing that some females had more full-brothers and/or fewer half-brothers given the genetic pedigree (table 2; fig. 4). This illustrates that patterns of extra-pair reproduction enacted by some female’s ancestors increased rather than decreased the number of possible matings between focal females and full-brothers versus half-brothers.

**Table 2:** Mean ±SD (and range) of the number of lifetime possible matings for individual adult female song sparrows at 15 specified relationships, and with more distant related and unrelated individual adult males, given the social and genetic pedigrees. The mean difference shows the mean decrease (negative values) or increase (positive values) in the number of possible matings at each relationship level given the genetic versus social pedigrees across 114 individual adult females. Full distributions of the differences are shown in fig. 4. *Z* and *p* denote the Wilcoxon rank sum test statistic value and associated *p* value. Relationships where numbers of possible matings decreased or increased significantly given the genetic pedigree are highlighted in bold.

| Relationship |  | Social pedigree | Genetic pedigree | Mean Difference | Z (p) |
| --- | --- | --- | --- | --- | --- |
| 1<sup>st</sup> degree | Father | 0.92 ±0.88<br>(0–5) | 0.85 ±0.88<br>(0–5) | -0.07 ±0.47<br>(-2–1) | 0.7<br>(0.49) |
|  | Full-brother | 1.50 ±1.86<br>(0–9) | 0.90 ±1.33<br>(0–6) | **-0.60 ±1.05**<br>**(-6–1)** | **2.9**<br>**(0.004)** |
|  | Son | 0.90 ±1.85<br>(0–11) | 0.90 ±1.85<br>(0–11) | 0.00 ±0.00<br>(0–0) | 0.0<br>(1.00) |
| 2<sup>nd</sup> degree | Grandfather | 0.40 ±0.74<br>(0–4) | 0.35 ±0.60<br>(0–2) | -0.05 ±0.65<br>(-4–2) | 0.2<br>(0.84) |
|  | Uncle | 0.78 ±1.17<br>(0–5) | 0.39 ±0.88<br>(0–5) | **-0.39 ±0.75**<br>**(-4–1)** | **3.4**<br>**(<0.001)** |
|  | Half-brother | 0.96 ±1.71<br>(0–10) | 2.54 ±2.87<br>(0–12) | **+1.59 ±2.17**<br>**(-1–11)** | **5.7**<br>**(<0.001)** |
|  | Double first cousin | 0.07 ±0.42<br>(0–3) | 0.00 ±0.00<br>-- | -0.07 ±0.42<br>(-3–0) | 2.0<br>(0.05) |
|  | Nephew | 1.39 ±2.97<br>(0–14) | 0.65 ±1.74<br>(0–12) | **-0.75 ±2.12**<br>**(-14–1)** | **2.2**<br>**(0.03)** |
|  | Grandson | 0.37±1.20<br>(0–7) | 0.39 ±1.48<br>(0–12) | +0.02 ±0.59<br>(-2–5) | 0.4<br>(0.69) |
| 3<sup>rd</sup> degree | Great-grandfather | 0.17 ±0.46<br>(0–2) | 0.15 ±0.55<br>(0–3) | -0.02 ±0.69<br>(-2–3) | 1.0<br>(0.32) |
|  | Single first cousin | 2.33 ±2.87<br>(0–16) | 0.76 ±1.77<br>(0–15) | **-1.57 ±2.44**<br>**(-13–2)** | **5.5**<br>**(<0.001)** |
|  | Great-grandson | 0.11 ±0.72<br>(0–7) | 0.12 ±0.81<br>(0–7) | +0.02 ±0.19<br>(0–2) | 0.0<br>(0.99) |
| 4<sup>th</sup> Degree | Half-uncle | 1.08 ±1.75<br>(0–8) | 2.00 ±2.30<br>(0–10) | **+0.92 ±2.09**<br>**(-6–6)** | **3.6**<br>**(<0.001)** |
|  | Half-single first cousin | 2.46 ±3.17<br>(0–17) | 5.39 ±6.44<br>(0–35) | **+2.94 ±5.53**<br>**(-13–28)** | **4.1**<br>**(<0.001)** |
|  | Half-nephew | 1.98 ±4.29<br>(0–23) | 2.82 ±5.46<br>(0–32) | +0.84 ±4.63<br>(-16–24) | 1.0<br>(0.33) |
| More distant |  | 47.92 ±35.06<br>(0–207) | 45.11±35.06<br>(0–182) | -2.81 ±7.65<br>(-29–20) | 0.6<br>(0.52) |
| Unrelated |  | 7.09 ±24.49<br>(1–261) | 7.09 ±24.49<br>(1–261) | 0.00 ±0.00<br>(0–0) | 0.0<br>(1.00) |

**Figure 4:**
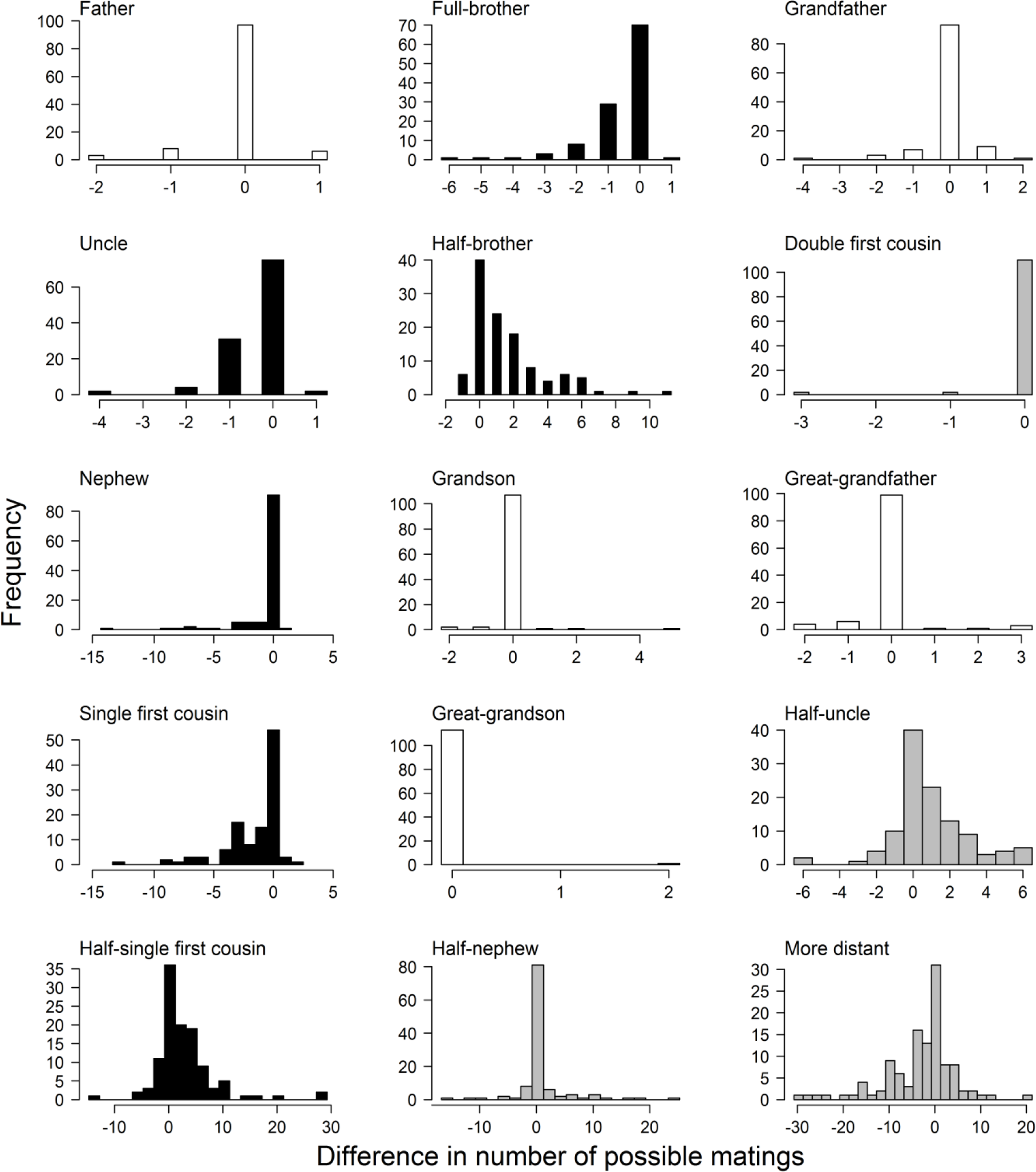
Distributions of the difference in the number of possible matings at each focal relationship level (listed in table 2) across 114 individual adult females given the genetic versus social pedigrees. Negative and positive values respectively indicate decreases and increases in the number of possible matings with available adult males at each relationship level. White bars denote lineal relatives (where little difference in the number of possible matings is expected), black bars denote relationship levels where the mean increase or decrease in the number of possible matings differed significantly from zero (table 2), and grey bars denote all other non-lineal relationship levels. Two relationship levels (‘son’ and ‘unrelated’) are not depicted because the difference in the number of matings between the social and genetic pedigrees was uniformly zero (table 2).

On average, there were also fewer possible matings between females and their full uncles, nephews, double first-cousins and single first-cousins given the genetic versus social pedigree and correspondingly increased numbers of possible matings with half-uncles and half-single first cousins (but little change in the number of possible matings with half-nephews, table 2). However, the distributions of the within-female differences again spanned zero, especially for half-single first cousins (fig. 4). There was consequently substantial among-individual variation in the consequences of extra-pair reproduction for the risk of inbreeding with 3^rd^ versus 4^th^ degree relatives.

As expected there was no change in the number of possible female-son matings given the genetic versus social pedigrees, and only small average changes in the numbers of possible matings with fathers, grandfathers, grandsons, great-grandfathers and great-grandsons (table 2) with little variation among individuals (fig. 4). Furthermore, there was little or no change in the number of possible matings between females and more distant relatives or completely unrelated males, respectively (table 2).

Overall, the individual-level differences in the distribution of relatives available as possible mates translated into substantial population-level differences: extra-pair reproduction meant that, across the population, adult females had 40% fewer possible matings with full-brothers, 166% more possible matings with half-brothers, and 85% more possible matings with 4^th^ degree relatives than with analogous 2^nd^ and 3^rd^ degree relatives (Supporting Information S4).

### Distribution of relatedness among possible mates

Due to variation in inbreeding among females’ ancestors, there was substantial among-individual variation in the mean kinship (*k*) between adult females and their possible mates that were identified as 1^st^, 2^nd^, 3^rd^ or 4^th^ degree relatives given the social pedigree (fig. 5A–D), particularly for 1^st^ and 2^nd^ degree relatives. Of the females that had ≥1 possible mate that was identified as a 1^st^, 2^nd^, 3^rd^ or 4^th^ degree relative given the social pedigree most, but not all, had lower mean *k* with these same sets of possible mates given the genetic pedigree (fig. 5A–D). Across females, mean *k_GEN_* was significantly lower than mean *k_SOC_* for all four categories of relative, but the magnitude of the difference was smallest for 4^th^ degree relatives (table 3). Conversely, mean *k_SOC_* and mean *k_GEN_* did not differ across females’ possible mates that were identified as more distant relatives given the social pedigree (table 3; fig. 5E). Because newly arrived immigrants were the only individuals that were completely unrelated to their possible mates, mean *k_SOC_* and mean *k_GEN_* were identical across individuals that were identified as non-relatives in the social pedigree (table 3; fig. 5F).

**Figure 5:**
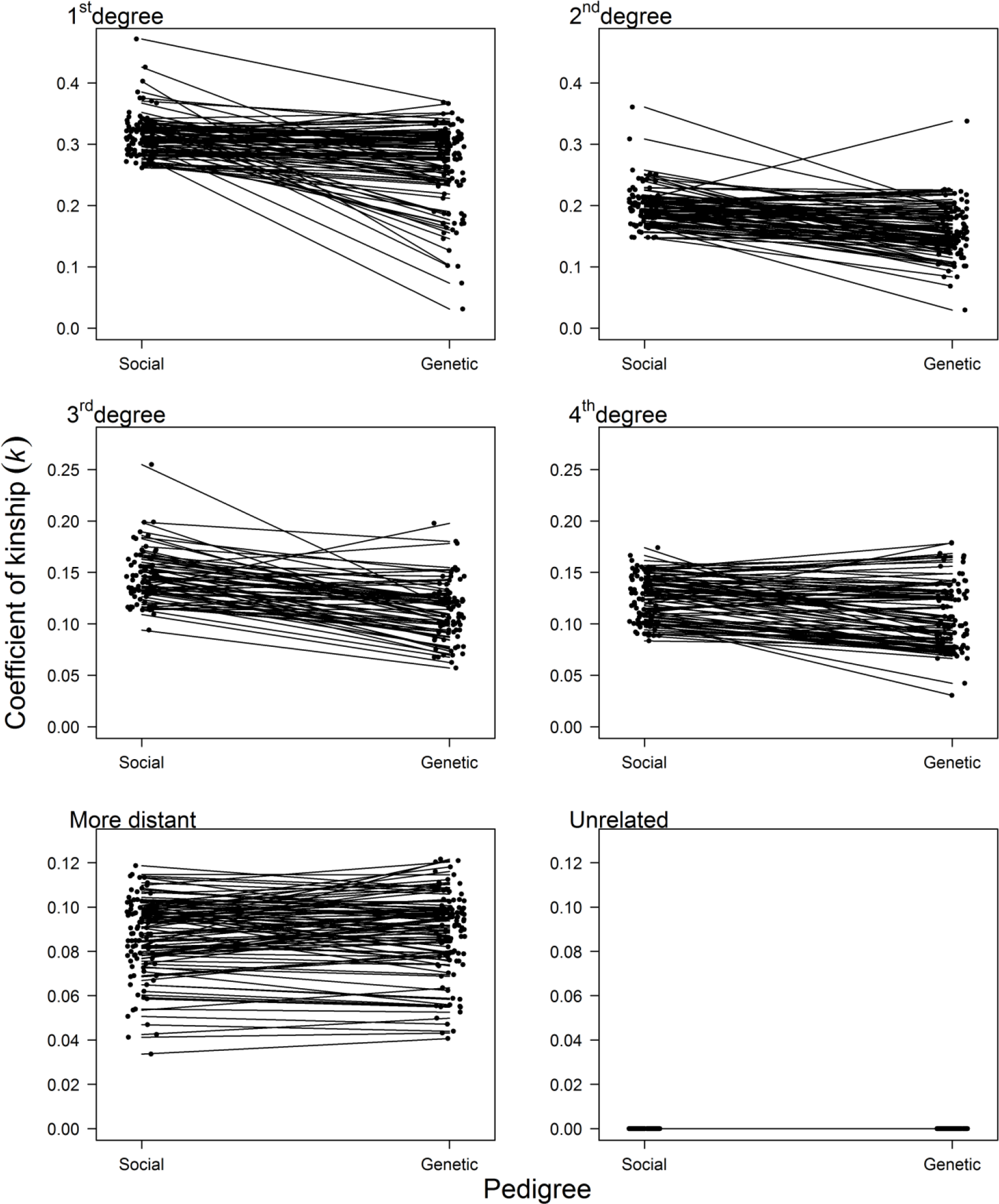
Mean coefficient of kinship (*k*) between individual adult female song sparrows and all possible adult male mates that were identified in the social pedigree as 1^st^, 2^nd^, 3^rd^ or 4^th^ degree relatives, or as more distant relatives or as unrelated, where *k* is calculated given the social pedigree (‘Social’) or genetic pedigree (‘Genetic’). Note that y-axis scales differ among rows of panels. Points denote individual females (jittered for clarity), and lines join observations for individual females given the two pedigrees. Of the females that had ≥1 possible mate that was identified as a 1^st^, 2^nd^, 3^rd^ or 4^th^ degree relative given the social pedigree most, but not all, had lower mean *k* with these same sets of possible mates given the genetic pedigree (80% [83/104], 84% [83/99], 87% [68/78] and 73% [71/97] of females respectively).

**Table 3:**
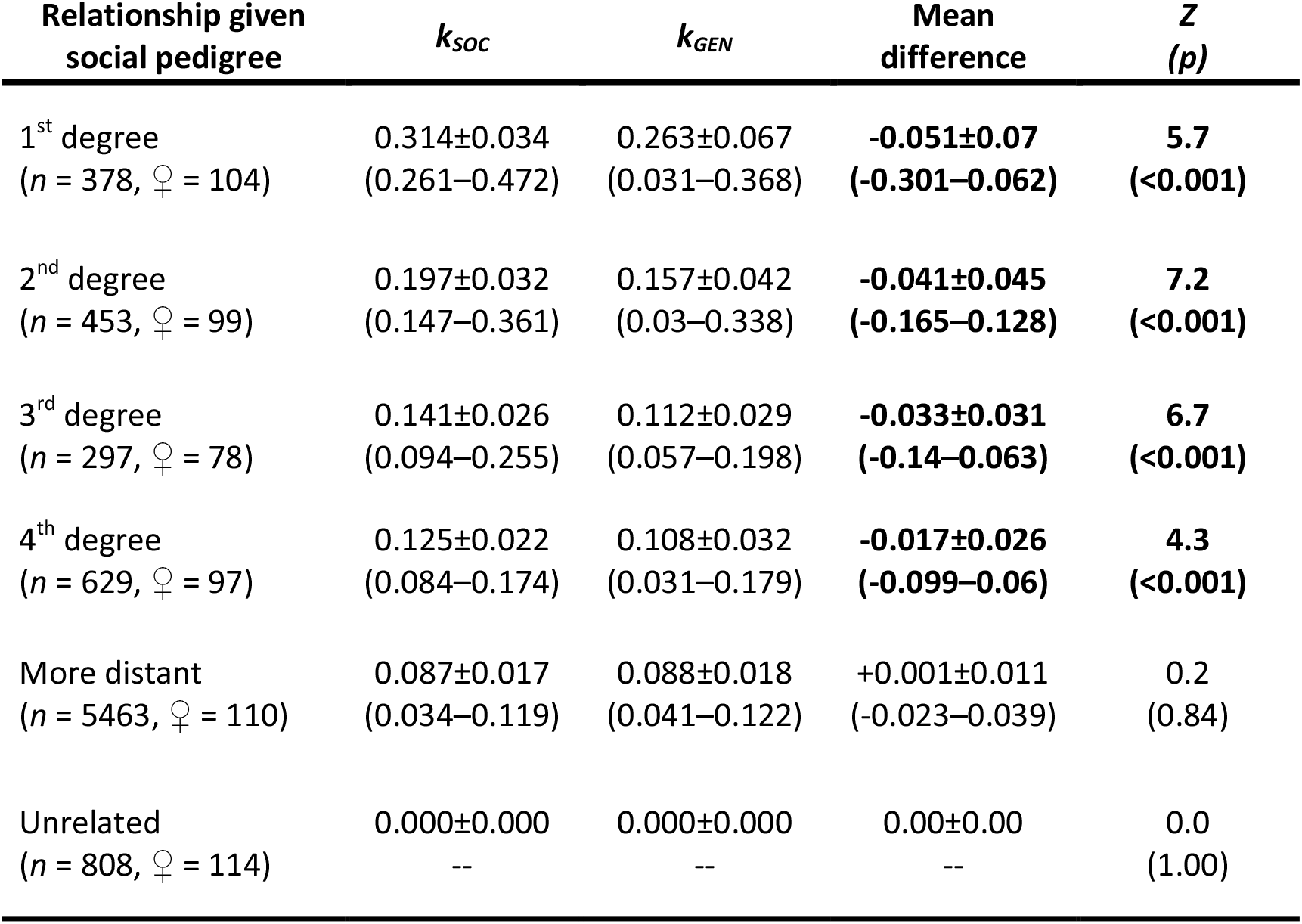
Mean ±SD (and range) pairwise coefficient of kinship (*k*) between individual adult female song sparrows and all possible adult male mates that were classified as 1^st^, 2^nd^, 3^rd^ or 4^th^ degree relatives, or as more distant relatives or as unrelated given the social pedigree, with *k* calculated from the social pedigree (*k_SOC_*) or genetic pedigree (*k_GEN_*). *n* and ♀ respectively represent the numbers of possible matings and individual females in each category. Mean difference denotes the mean decrease (negative values) or increase (positive values) in mean *k* for individual females given the genetic versus social pedigrees (i.e., *k_GEN_ - k_SOC_*). *Z* and *p* denote the Wilcoxon rank sum test statistic value and associated *p* value. Degrees of relationship where mean *k* decreased significantly are highlighted in bold.

Of the 114 females, 71% (81) had negative values of mean *k_DIFF_* across all possible matings given the genetic pedigree versus the social pedigree, while 25% (29) had positive values of mean *k_DIFF_*, and 4% (4) had no change in mean *k_DIFF_* (three female immigrants that were alive in only one year, and one female immigrant whose only possible matings with relatives were with sons or grandsons). Grand mean *k_DIFF_* across all possible matings for individual females was -0.007 ±0.01 (median -0.008, range -0.035–0.017), showing that, on average, females were slightly less related to all possible mates given the genetic pedigree than given the social pedigree.

However, across all pooled possible matings for all females, the distributions of *k_GEN_* and *k_SOC_* were significantly different (two-sample Anderson-Darling test, *AD* = 28.27, *T* = 35.79, *p* < 0.001). This difference arose because the distribution of *k_GEN_* included fewer possible matings at higher *k* (fig. 6, black bars), but more possible matings at lower but nonzero *k* (fig. 6, white bars), than the distribution of *k_SOC_*. There was again no difference in the number of possible matings among unrelated individuals (i.e., *k* = 0, fig. 6). Thus, the main effects of extra-pair mating were not in altering mean relatedness among potential mates but in altering the distribution of relatedness, such that females were less likely to mate at intermediate and higher levels of *k* (i.e., with closely related males) and more likely to mate at lower, but non-zero, levels of *k* (i.e., with more distantly related males).

**Figure 6:**
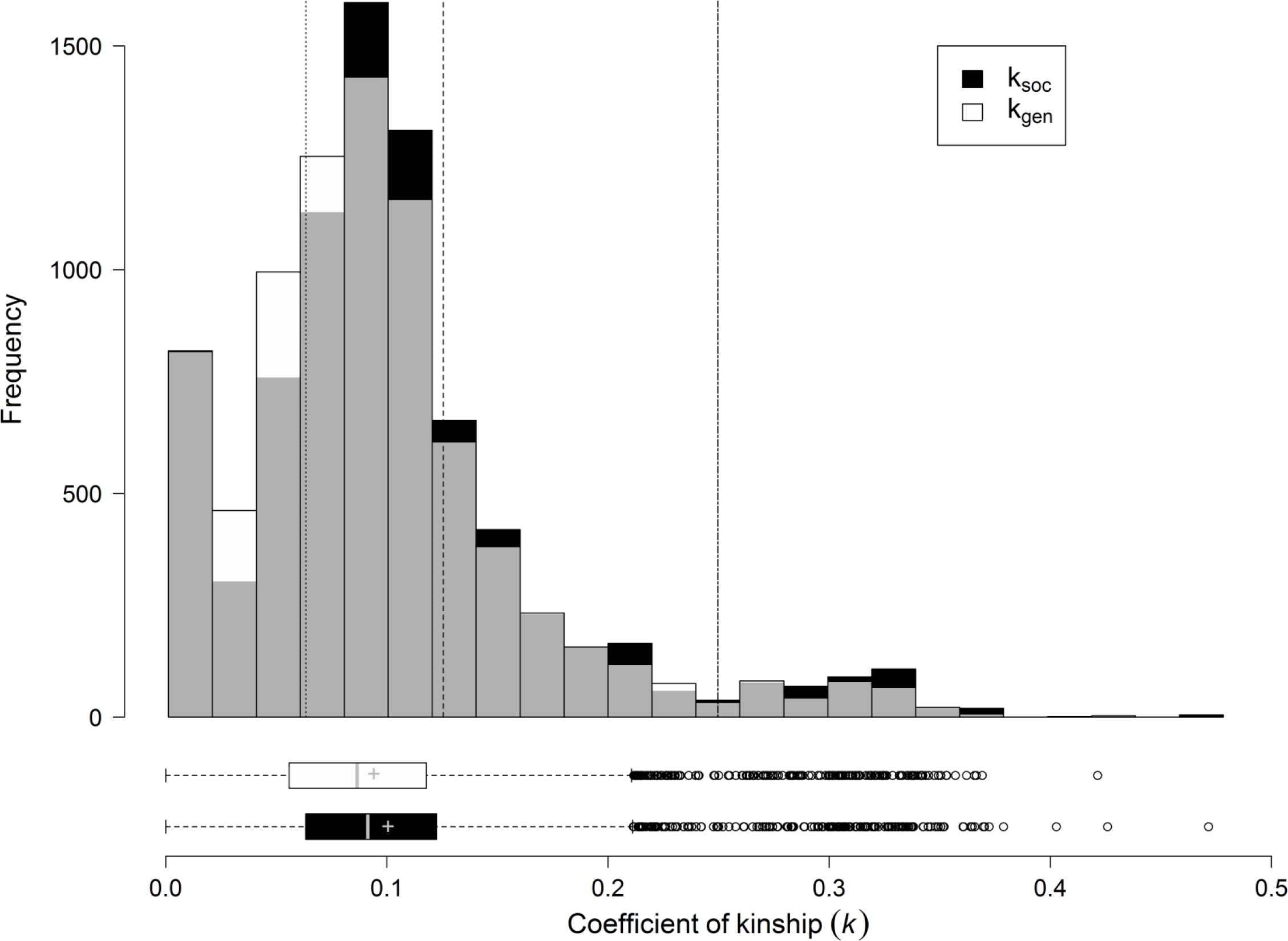
Overall distributions of pairwise coefficients of kinship across all possible matings between adult female and male song sparrows, calculated from the social pedigree (black bars) or genetic pedigree (white bars), with grey bars denoting overlap between the two distributions. Dotted, dashed, and dot-dashed lines depict kinship values equivalent to first cousin (0.0625), half-sib (0.125), and full-sib (0.25) matings, respectively. Box plots further visualize the distribution of *k* given each pedigree, where box lines represent the median, upper and lower quartiles, whiskers demarcate 1.5× the interquartile range, and ‘+’ represents the mean. Mean *k_SOC_* was 0.101±0.069 (median 0.914, range 0.000–0.472) and mean *k_GEN_* was 0.094±0.065 (median 0.087, range 0.00–0.421), corresponding to a small but statistically significant mean decrease of -0.006±0.040 (median -0.003, range -0.301–0.251, Wilcoxon signed rank test: *Z* = 18.95, *p* < 0.001).

## Discussion

Simultaneous polyandry is widely hypothesized to have evolved to facilitate inbreeding avoidance in populations where relatives interact and inbreeding depression is strong (e.g., Stockley et al. 1993; Tregenza and Wedell 2000; Michalczyk et al. 2011; Duthie et al. 2016; Bocedi and Reid 2017). Consequently, numerous empirical studies on diverse systems have tested whether polyandrous females avoid inbreeding by expressing pre-copulatory and/or post-copulatory choice for less closely related mates (Tregenza and Wedell 2002; Firman and Simmons 2008; Brouwer et al. 2011; Reid et al. 2015*a*). However, no studies have quantified the degree to which intrinsic effects of polyandry on sibship structures might indirectly reduce inbreeding risk (i.e., the ‘indirect inbreeding avoidance hypothesis’, IIAH) in systems experiencing natural variation in polyandry, polygyny and paternity within and across overlapping generations. Accordingly, we compared long-term social and genetic pedigree data from free-living song sparrows to examine the consequences of extra-pair paternity, and hence of underlying simultaneous polyandry, for sibship structures and resulting distributions of relationships and relatedness. Further, by comparing observed patterns to those that would have arisen given lifelong monandry (i.e., no simultaneous or sequential polyandry) or given simultaneous polyandry but no resulting polygyny (i.e., the ‘distinct males assumption’), we isolated effects of major components of the complex overall natural mating system on sibship structures.

Comparisons between social and genetic pedigrees have previously been used to quantify effects of extra-pair reproduction on the variance in male reproductive success and hence on effective population size and the opportunity for selection (Webster et al. 1995, 2007; Freeman-Gallant et al. 2005; Lebigre et al. 2012). Such effects are often small, including in song sparrows (Lebigre et al. 2012, see also Karl 2008). However, such results do not preclude the possibility that extra-pair reproduction could affect individual-level inbreeding risk. This is because the same overall variance in male reproductive success, but very different sibship structures and distributions of relationships and relatedness, can arise if individual males sire several offspring of one female (i.e., generating full-sibs) or sire one offspring of several females (i.e., generating paternal half-sibs).

### Sibship structures

It may seem inevitable that extra-pair reproduction will reduce full-sibships, as assumed by the basic IIAH, and by Cornell and Tregenza’s (2007) initial theoretical model. However our analyses illustrate that such effects arising within a natural mating system are not so straightforward. Comparison of the social and genetic song sparrow pedigrees showed that extra-pair reproduction did indeed increase the mean number of different males that sired individual females’ offspring and hence reduce the mean proportion of full-sibships (Prop_Full-sibs_) and increase the mean proportion of maternal half-sibships (Prop_Half-sibs_) among females’ lifetime banded offspring. However, such means mask substantial among-female variation, including cases where extra-pair reproduction increased rather than reduced Prop_Full-sibs_ (fig. 2B). Such patterns can result from non-independent extra-pair paternity when females produce numerous extra-pair offspring with the same male across broods (as indicated by fig. 3A,C), and/or if a female’s extra-pair male from one brood becomes her socially-paired male for another brood (or vice versa). Further, comparisons with the hypothetical occurrence of lifelong monogamy showed that the occurrence of social re-pairing across breeding attempts (i.e., sequential polyandry) already reduced the effects of simultaneous polyandry on sibship structures by ~26%. Selection for simultaneous polyandry stemming from the IIAH process might consequently be weaker given iteroparity and associated repairing than given semelparity and/or strict lifelong monogamy. Comparison with the hypothetical ‘distinct males’ pedigree showed that 68% of females would have had lower Prop_Full-sibs_ among their banded offspring in the absence of polygyny than given the observed pattern of polygyny defined by the genetic pedigree (fig. 2B). This implies that Cornell and Tregenza’s (2007) theoretical formulation of the IIAH might overestimate indirect selection on polyandry arising in polygynandrous systems.

While simultaneous polyandry can clearly affect the sibship structure of females’ conceived offspring, its consequences for inbreeding risk (and other kin interactions including kin cooperation and competition) ultimately depend on its effects on the sibship structure of offspring that survive to life-history stages when key interactions occur. In song sparrows, further comparisons of the genetic and social pedigrees showed that the effects of extra-pair reproduction on sibship structures were qualitatively similar, but subtly different, across recruited versus banded offspring (fig. 2). Most notably, Prop_Full-sibs_ for recruits no longer differed between the genetic and ‘distinct males’ pedigrees (table 1B,D vs F). These patterns imply that theoretical predictions regarding indirect selection on polyandry might, in some instances, be relatively robust to an assumption of no polygyny. However, such inferences from observed genetic and social pedigrees require the additional, and commonly violated, assumption that offspring survival to recruitment does not depend on paternity. In song sparrows, female extra-pair offspring are less likely to recruit than female within-pair offspring reared in the same brood (i.e., maternal half-sisters, Sardell et al. 2011), and extra-pair offspring of both sexes have lower survival and/or reproductive success than within-pair offspring in other passerine birds (e.g., house sparrows, *Passer domesticus*, Hsu et al. 2014; coal tit, *Periparus ater*, Schmoll et al. 2009). Any small reduction in inbreeding among polyandrous females’ offspring might therefore be further reduced by stochastic and/or deterministic variation in survival of offspring sired by different males. The ultimate consequences of polyandry for the expected frequency of close inbreeding and consequent fitness among descendants of polyandrous females in natural populations may therefore be smaller than predicted by models that do not consider differential offspring survival (e.g., Cornell and Tregenza 2007), and estimated in laboratory populations where variation in survival may be minimized (e.g., Power and Holman 2014). Future theoretical and empirical studies considering the evolutionary causes and consequences of polyandry arising through its effects on sibship structures should therefore consider such effects within the context of the overall mating system, including natural variation in paternity arising through sequences of polygyny and mate fidelity, re-pairing due to divorce and mate death, as well as differential offspring survival.

### Distributions of relationships and relatedness

The effects of simultaneous polyandry on sibship structures among recruited offspring are likely to alter the frequencies of diverse types of half-relatives versus full-relatives spanning multiple (overlapping) generations, thereby altering any individual’s overall potential for inbreeding or interacting with different types of relatives. The form and magnitude of indirect selection on polyandry stemming from the IIAH process might then differ from that predicted in restricted situations with within-brood mating and non-overlapping generations (e.g., Cornell and Tregenza 2007). Indeed, our comparisons of the social and genetic pedigrees of female and male song sparrows that survived to adulthood showed that ancestral extra-pair reproduction generally reduced the potential for inbreeding among different degrees of full-relatives, and increased the potential for inbreeding among more distant half-relatives. However, this change was not consistent across all individual females and types of relationship (table 2, fig. 4, Supporting Information S4). Similarly, simultaneous polyandry reduced the mean kinship (*k*) between adult females and their possible mates, most notably with available adult males that would otherwise have been 1^st^ degree relatives (fig. 5). However, the overall conclusions remained unchanged when all possible matings among 1^st^ degree relatives were excluded (Supporting Information S5), thereby considering a scenario where individuals actively avoid inbreeding with 1^st^ degree relatives, as could be achieved through some form of active or passive kin discrimination (e.g., Stow and Sunnucks 2004; Gerlach and Lysiak 2006; Archie et al. 2007; Brouwer et al. 2011; Ihle and Forstmeier 2013). Overall, the individual-level differences in relatedness among possible mates stemming from simultaneous polyandry resulted in fewer possible matings at intermediate and higher *k* (i.e., among closely related pairs), and more possible matings at lower but nonzero *k* (fig. 6).

Such conclusions rely on the implicit assumptions of our study design that mating decisions and recruitment are unaffected by pedigree structure, and hence that there is no active inbreeding avoidance or differential survival by within-pair versus extra-pair offspring. Indeed, previous analyses showed that song sparrows do not actively avoid inbreeding through social pairing or extra-pair reproduction (Keller and Arcese 1998; Reid et al. 2015*a*). However, to further consider the implications of such assumptions, we conducted additional analyses to quantify effects of polyandry on relatedness within a single cohort (Supporting Information S6). Such analyses have the advantage that they do not require any assumptions regarding patterns of mating or survival in the absence of extra-pair reproduction, but the disadvantage that they eliminate effects of polyandry on relatedness generated across multiple (overlapping) generations. These analyses also showed reduced potential for close inbreeding (*k* ≥ 0.25) given the genetic versus social pedigrees, but no reduction in more distant inbreeding (0.03125 ≤ *k* < 0.25, Supporting Information S6). These supporting results illustrate that overall effects of polyandry in reducing the potential for inbreeding at intermediate *k* accumulate across generations, meaning that exact quantitative outcomes could be influenced by patterns of differential survival of within-pair versus extra-pair offspring.

### Implications

Our results imply that the magnitude and direction of indirect selection on simultaneous polyandry stemming from the intrinsic consequences of such polyandry for distributions of *k* among females’ offspring, and hence grand-offspring *f*, will depend on the shape of the relationship between fitness and *f* (i.e., the form of inbreeding depression). Given multiplicative effects of deleterious recessive alleles, inbreeding depression is expected to be log-linear, such that the reduction in fitness decreases with increasing *f* (fig. 7, Morton et al. 1956; Charlesworth and Charlesworth 1987; Charlesworth and Willis 2009). Counter-intuitively, under these conditions, polyandry might in fact cause a net decrease in mean fitness, even though it slightly reduces mean grand-offspring *f*. Intrinsic indirect selection on polyandry stemming from ‘indirect inbreeding avoidance’ might then impede rather than facilitate polyandry evolution. However, given epistatic or threshold effects, inbreeding depression could be weak up to some value of *f* above which fitness decreases markedly (e.g., fig. 7, Charlesworth and Willis 2009). Given such threshold effects, the long-term relative frequency of alleles underlying polyandry could then increase due to the reduced frequency of matings among close relatives and the resulting net increase in mean offspring fitness that would arise despite an increased frequency of matings among more distant relatives.

**Figure 7:**
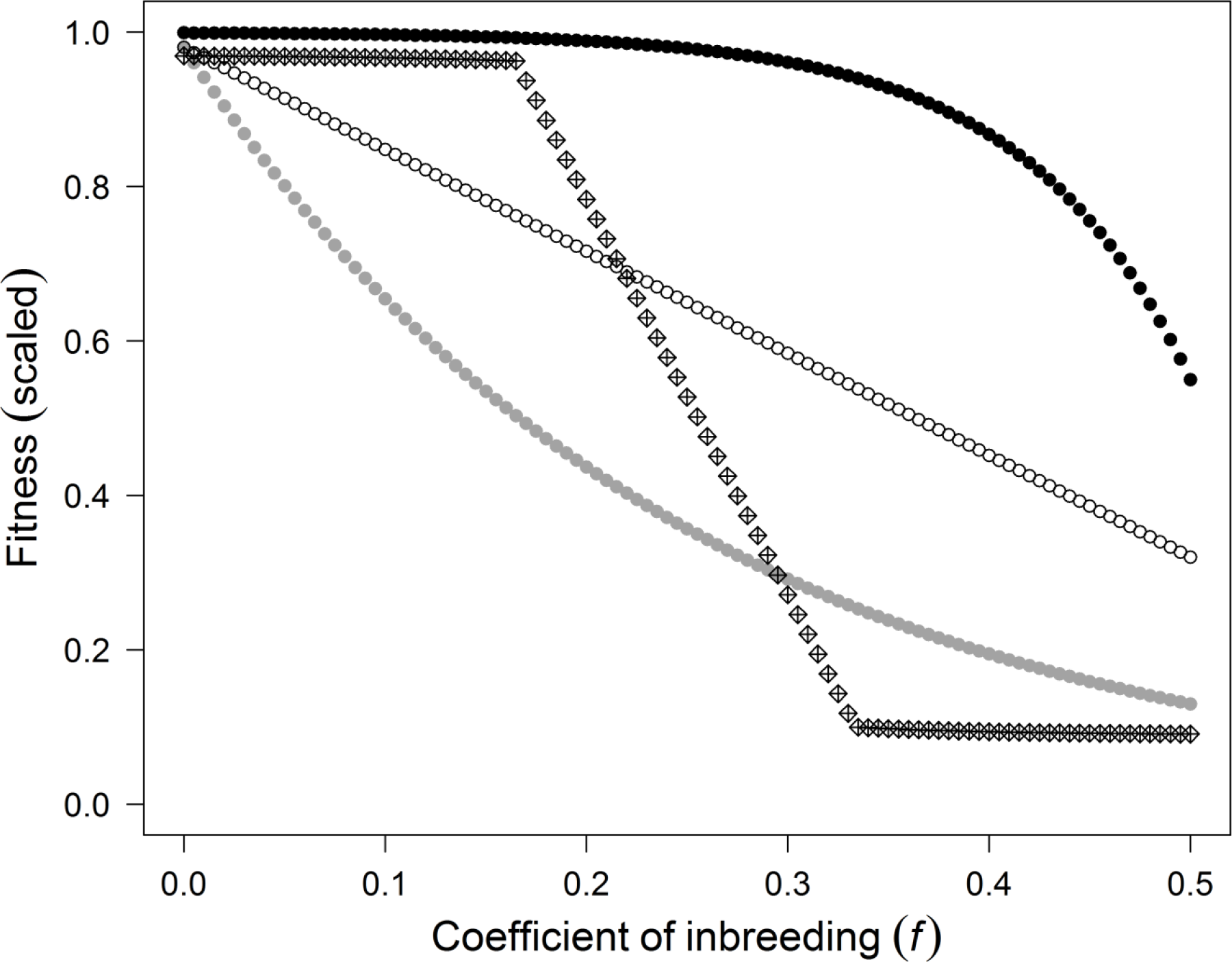
Conceptualization of potential relationships between mean fitness (scaled relative to an outbred individual) and individual coefficient of inbreeding (*f*). Such relationships could be linear (white circles), concave (i.e., log-linear, given multiplicative allelic effects, grey circles), convex (given epistasis, black circles), or follow a threshold pattern (diamonds). Points denote x-axis intervals of 0.01 to depict the effects of different shapes of inbreeding depression on equal scales. Note that concave and convex forms of inbreeding depression are often depicted on a log scale (i.e., log-fitness) such that log-linear effects appear linear (e.g., Charlesworth and Willis 2009). Each series of points is meant to convey qualitative patterns of inbreeding depression and not quantitative values, and so are jittered for clarity.

The form of inbreeding depression is very difficult to quantify in natural populations, not least because close inbreeding often occurs infrequently and may be more likely in high-fitness lineages where more relatives are available for mating, meaning that phenotypic effects of inbreeding could be confounded with environmental and/or additive genetic effects (Reid et al. 2008). Experimental assessments of the shape of inbreeding depression across ranges of *f* relevant to animal mating systems are also scarce, because most experimental studies consider restricted inbred groups generated through one or multiple generations of sib-sib mating (Charlesworth and Charlesworth 1987; Keller and Waller 2002; Charlesworth and Willis 2009). Full quantitative, mechanistic evaluation of the ‘indirect inbreeding avoidance’ process in driving or impeding polyandry evolution will therefore require information on distributions of sibships, relationships and relatedness arising within complex natural mating systems to be coupled with detailed experimental assessments of the form of inbreeding depression arising across appropriate ranges of *f*.

## Acknowledgments

We thank the Tsawout and Tseycum First Nations for access to Mandarte Island, Pirmin Nietlisbach, Lukas Keller, Greta Bocedi, Brad Duthie, and Matthew Wolak for helpful discussions, and the European Research Council, National Sciences and Engineering Research Council of Canada, and Swiss National Science Foundation for funding. Field data collected following UBC Animal Care Committee (A07-0309) and Environment Canada (Master banding permit 10596) guidelines. All data from this publication will be archived in the Dryad Digital Repository (*doi* upon acceptance).

## Supplementary Material

**Supporting Information S1:** Quantifying the combined effects of simultaneous and sequential polyandry on sibship structures

**Supporting Information S2:** Sibship structure among banded offspring of females that produced recruits

**Supporting Information S3:** Sibship structure among males’ banded and recruited offspring

**Supporting Information S4:** Population-wide effects of polyandry on the distribution of relatedness

**Supporting Information S5:** Distribution of relatedness excluding all 1^st^ degree relatives

**Supporting Information S6:** Distribution of relatedness within cohorts

